# Myelin Imaging Using Dual-echo 3D Ultra-short Echo Time MRI with Rosette k-Space Pattern

**DOI:** 10.1101/2021.09.18.460869

**Authors:** Xin Shen, Ali Caglar Özen, Antonia Sunjar, Serhat Ilbey, Riyi Shi, Mark Chiew, Uzay Emir

**Author notes:** Corresponding author Uzay Emir, Purdue University, School of Health Sciences, 550 Stadium Mall Drive, West Lafayette, IN 47907-2051, Institution information, Purdue University, 610 Purdue Mall, West Lafayette, IN, USA, 47907.

## Abstract

**Purpose:** This study aimed to develop a new 3D dual-echo rosette k-space trajectory, specifically for ultra-short echo time (UTE) magnetic resonance imaging (MRI) applications. The direct imaging of the myelin bilayer, which has ultra-short transverse relaxation time (uT_2_), was acquired to test the performance of the proposed UTE sequence.

**Theory and Methods:** The rosette trajectory was developed based on rotations of a ‘petal-like’ pattern in the k_x_-k_y_ plane, with oscillated extensions in k_z_-direction for 3D coverage. Five healthy volunteers were recruited and underwent ten dual-echo 3D rosette UTE scans with various echo times (TEs). Dual-exponential complex model fitting was performed on the magnitude data to separate uT_2_ signals, with the output of uT_2_ fraction, uT_2_ value, and long T_2_ value.

**Results:** The reconstructed images’ signal contrast between whiate matter (WM) and grey matter (GM) increased with longer TEs. The WM regions had higher uT_2_ fraction values than GM (10.9%±1.9% vs. 5.7%±2.4%). The uT_2_ value was approximately 0.15 milliseconds in WM.

**Conclusion:** The higher uT_2_ fraction value in WM compared to GM demonstrated the ability of the proposed sequence to capture rapidly decaying signals.

## Introduction

Conventional ^1^H-Magnetic Resonance Imaging (MRI) sequences are widely used for *in vivo* imaging, which focus on detecting signals from tissues with relatively long spin-spin (transverse) relaxation time (T_2_). However, depending on the surrounding chemical environment, some protons in specific tissues in the human body have an ultra-short T_2_, which conventional MRI can hardly detect due to the relatively long echo time (TE) in the order of milliseconds (ms). For example, the myelin bilayer in cerebral white matter (WM) (1), cortical and trabecular bones (2), ligaments (3), tissues with high iron concentration (4) have ultra-short T_2_ values. On the other hand, ultra-short echo time (UTE) MRI (5) sequences are capable of acquiring data with TE in the order of microseconds (μs), which can provide images of tissues with ultra-short T_2_ directly.

For UTE sequences, the readout gradients are applied immediately after the completion of the RF pulses. Therefore, to achieve minimum possible TE, each data acquisition in UTE sequences needs to follow a center-out trajectory. Although 2D UTE methods are available, they have limitations making it difficult to achieve an appropriate slice selection and a minimum TE due to eddy currents and imperfect gradients (6,7). Alternatively, the most common k-space trajectory used in UTE MRI applications is a 3D radial center-out readout. However, sampling the outer k-space with a radial pattern may not be efficient (8). Therefore, novel 3D k-space trajectories with greater curvature per spoke have been proposed for a more efficient sampling strategy, e.g., the spiral-like extended cones sampling (7,8). Other state-of-the-art 3D UTE techniques include density-adapted 3D radial UTE (9), FLORET UTE sequence (10) and a 3D *koosh ball* trajectory UTE sequence (11). Rosette k-space trajectories, which allow a center-out sampling pattern while providing more samples in the outer k-space per spoke than radial trajectories, are also potential candidates for 3D UTE MRI. In addition, the rosette k-space trajectory samples data in a more incoherent pattern than the radial trajectory. Therefore, it offers the potential for further acceleration using higher under-sampling factors and the compressed sensing technique for reconstruction (12). However, 3D rosette k-space patterns have not yet been demonstrated in UTE applications.

Alternatively, a similar MRI-based technique, zero echo time (ZTE) MRI, has also been proposed for imaging ultra-short T_2_ tissue. However, unlike UTE imaging, the readout gradients are applied before the RF pulse and are always on in ZTE sequences (13). In addition, the center k-space in the ZTE sequence is empty because of the dead time due to the switching from RF pulse excitation to data receiving (13), which requires additional scans or interpolation based on an over-sampling strategy to fill in the missing k-space (14). Nevertheless, a recent comparison study has shown that UTE and ZTE MRI sequences provide similar results in terms of image quality and signal-to-noise ratio (SNR) (15).

One of the essential applications of 3D UTE and ZTE sequences is imaging the myelin bilayer in brain white matter (WM). Myelin constituents are water and dry mass, in which the dry mass is composed of about 80% lipids and 20% protein (16). Since transverse relaxation of the lipid proton signal becomes shorter in this geometrically restricted environment, T_2_ values vary from a few microseconds to milliseconds, where 75% of the myelin lipid signal manifests T_2_ values below 100 *μs* (17). Thus, conventional MRI sequences with TEs in the order of milliseconds or longer can hardly capture the rapid signal decay of the lipid bilayer.

Many alternative MRI-based methods have been proposed for myelin imaging, e.g., the methods based on magnetization transfer (MT) effects, separation of myelin water signals, susceptibility imaging, and cortical myelin mapping (18-21). However, while those methods provide an indirect measurement of myelin (22), UTE MRI can provide a direct measure of myelin, which may be more specific and more precise (7).

UTE MRI methods have shown great promise for imaging the signals arising from the phospholipid chains of myelin sheaths. Common techniques for the separation of myelin signals based on UTE acquisitions are the combination of inversion recovery (IR) suppression (1) and dual-echo subtraction (23). Even though those methods showed the potential to suppress signals originating from long-T_2_ components in both grey matter (GM) and WM, the inversion time (TI) of IR suppression is difficult to choose because of the wide range of longitudinal relaxation time (T_1_) reported in WM (700 to 1100 ms at 3T) (24,25). An alternative method to quantify myelin signals was using a dual-exponential model fitting with data input based on a multiple TEs acquisition, ranging from microseconds to milliseconds.

This study aims to develop a novel rosette k-space pattern for 3D UTE MRI and test the sequence by directly and non-invasively measuring the myelin bilayer with whole-brain coverage. With the novel k-space design, dual-echo data is sampled in a single acquisition, allowing an extended TE coverage for the dual-exponential complex model fitting. In addition, compressed sensing and low-rank denoising were applied to reconstruct images from the acquired non-Cartesian k-space data.

### Theory

Rosettes are non-Cartesian k-space trajectories with high design flexibility (26). The shape of the rosette trajectory depends highly on the parameters, which could be identical to rings or radial patterns in extreme cases. The rosette trajectories are well known for the multiple crossings of k-space origin (27), which suggests the potential for multiple-echo acquisition.

The following equations define the 3D rosette k-space trajectory (28):

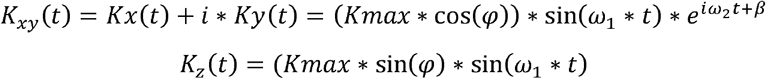

where *Kmax* is the maximum extent of k-space, *ω*_l_ is the frequency of oscillation in the radial direction, *ω*_2_ is the frequency of rotation in the angular direction, *φ* determines the location in the z-axis, and *β* determines the initial phase in the angular direction.

In Figure 1A, a rosette trajectory is shown for the specific case where *ω*_l_ and *ω*_2_ are set to be equal. Origin of the k-space is sampled at the beginning and the end of each repetition time (TR), forming a petal-like sampling pattern (Figure 1A). Dual-echo images can be generated within a single acquisition with a manual separation at the middle of each data readout (Figure 1A and 1B). TE values are separately determined by the time of two crossings of the k-space origin. The gradients of one petal trajectory are shown in Figure 1B. The amplitude of the readout gradient began at zero to avoid any potential delays caused by the gradients’ ramp-up (details in discussion).

**Figure 1.**
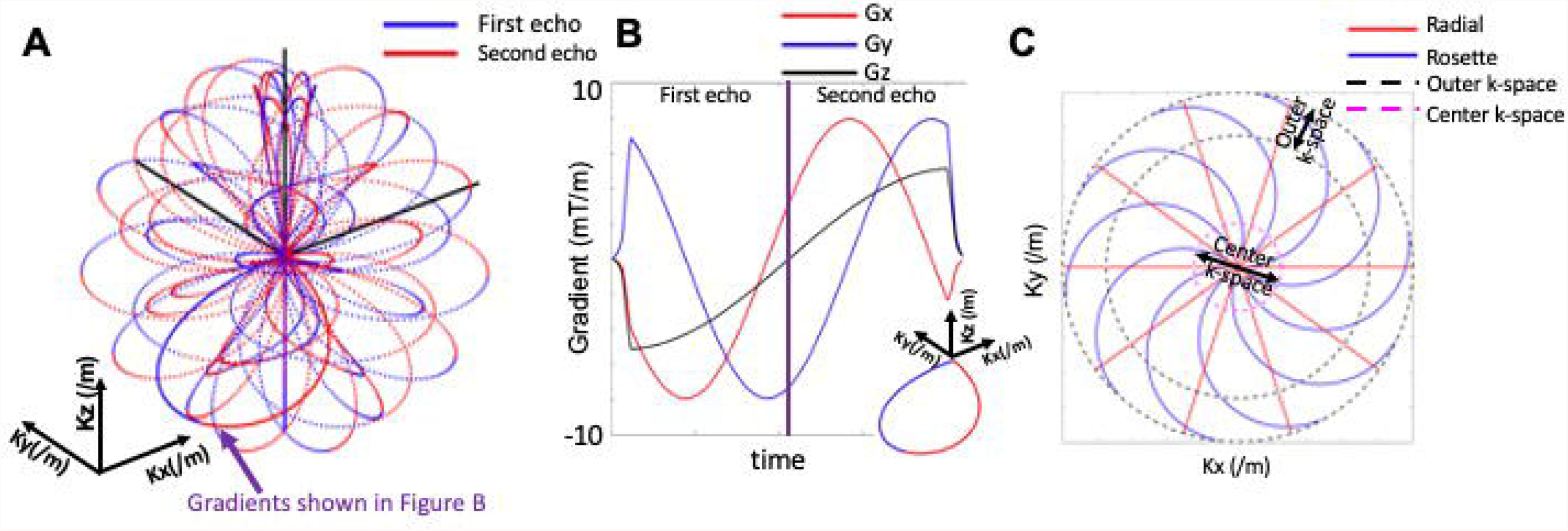
Illustrations of the 3D rosette k-space design. A) Selective spokes with varied rotations in the k_x_-k_y_ plane and varied extensions in the k_z_-axis are shown. Each spoke crosses the origin of k-space at the beginning and the end, forming a ‘petal-like’ pattern. The dual-echo acquisition (blue: first echo, red: second echo) was generated based on manual separation at the middle of each ‘petal-like’ pattern. B) The gradients in all three directions (red: gradients in the x-axis, Gx; blue: gradients in the y-axis, Gy; black: gradients in the z-axis, Gz) of a selective spoke as shown in Figure 1A. All gradients began at zero amplitude to avoid any potential delays caused by the gradients’ ramp-up. C) A 2D example to illustrate the differences between rosette (blue) and radial (red) acquisitions in peripheral (outer) k-space coverage (regions between black dash lines). With greater curvature, rosette trajectory provides more samples in the peripheral k-space than radial trajectory.

The parameters for the UTE acquisition were: *Kmax*=250/m, *ω*_l_=*ω*_2_=1.611 kHz, number of total petals= 36100, samples per petal=210, *φ* was sampled uniformly in the range of [−*π*/2, *π*/2], and *β* was sampled uniformly in the range of [0,2*π*]. With ten *μs* readout dwell time, the difference between first TE (TE_1_) and second TE (TE_2_) is equal to 210 × 10 *μs* = 2.1 *ms*, (details of other scan parameters are stated in the Methods section). Based on Nyquist criteria calculation, to avoid artifacts, the minimum number of total petals should be 4*π* * *r*^2^ ≈ 45239, where r represents half of the 1D matrix size (r=60 in this paper). Due to the 3D rosette k-space pattern (i.e., the curvature and k-space coverage), this 80% (36100/45000) undersampling pattern did not suffer significant artifacts. The noise generated by the undersampling strategy was primarily removed by compressed sensing reconstruction. Because of these reasons, the trajectory with 36100 petals is considered as a full k-space pattern in this study. The Nyquist criteria for samples per petal varied depending on *φ* and *β*. It reaches the maximum required samples when *φ* = 0, which is *π* * *r* = 189 samples per petal (the number is for two TEs). Crusher gradients in all three directions were applied at the end of each readout gradient (not shown in Figure 1B).

The 3D radius (including all x, y, and z directions) of the rosette trajectory is expressed with a sine function (*Kmax* * sin(*ω*_l_ * *t*)), which varies within [0, *Kmax*]. In addition, the 3D radius of the rosette trajectory has a fast rate of change at small radii of k-space but a slow velocity at large radii of k-space. With the constant readout dwell time, the k-space velocity difference led to increased samples with the radii of k-space for each for each readout spoke (about 43% samples at radii of k-space larger than 0.75 *Kmax*). However, readout spoke, the 3D radius of the standard radial trajectory has a constant change rate, which led to a uniform sampling with the 1D radii of k-space (25% samples at radii of k-space larger than 0.75*Kmax*). A 2D example is illustrated in Figure 1C.

## Methods

The study was approved by the Institutional Review Boards (IRBs) of Purdue University. Five healthy subjects underwent brain scans with a whole-body 3T MRI scanner (Siemens Healthineers, Erlangen, Germany). A vendor-supplied 20-channel receiver head coil was used for all volunteers. An MPRAGE sequence was performed before the UTE sequence for the anatomical reference. The parameters used in the dual-echo 3D UTE sequence with rosette k-space sampling were: field of view (FOV)=240×240×240 mm^3^, matrix size=120×120×120, readout dwell time=10 *μs*, flip angle=7-degree, TR=7 ms, readout duration=2.1 ms, and RF pulse duration=10 *μs*. Ten repeated dual-echo UTE scans were performed with varied TEs. The first TEs were 20 *μs*, 40 *μs*, 60 *μs*, 80 *μs*, 100 *μs*, 150 *μs*, 300 *μs*, 600 *μs*, 1ms, and 1.5 ms. The second TEs were 2.12, 2.14, 2.16, 2.18, 2.2, 2.25, 2.4, 2.7, 3.1, and 3.6 ms. For one dual-echo UTE acquisition, 36100 petals were acquired (full k-space acquisition without acceleration factor), resulting in a scan duration of 4.2 minutes (36100×7 ms). The ten repeated UTE scans lasted 42 minutes, and the total scan time was about 50 minutes for each volunteer.

Image reconstruction and post-processing steps were performed in MATLAB (MathWorks, USA). Nonuniform fast Fourier transform (NUFFT) was used to calculate the forward encoding transform of the acquired k-space data (29). A compressed sensing approach was used for image reconstruction, using total generalized variation (TGV) as the sparsifying penalty (30). The complex-valued coil sensitivity maps were first extracted from the center of k-space using ESPIRIT for coil combination (31). Then, the reconstructed complex image data was phase-corrected for each echo time and resulted in the magnitude image data.

The post-processing procedure was performed using FSL (FMRIB Software Library) and SPM (Statistical Parametric Mapping) software. The workflow of post-processing steps was summarized in Figure 2, including registration to T1-weighted anatomy images (32,33), brain skin and skull removal (34), bias-field correction (35), voxel-wised LORA (low-rank approximations) correction (36), and voxel-wised dual-exponential fitting based on the magnitude of the free induction decay (FID) signal. Ten dual-echo UTE acquisitions allow to utilize additional 10 second TEs in the model for dual-exponential fitting. The fitting model was expressed as the equation (37,38):

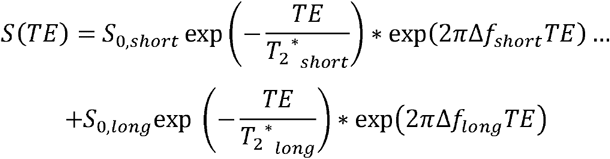

where *s*_0,*short*_ and *s*_0,*long*_ are the proton density, and Δ*f*_*short*_ and Δ*f*_*long*_ are the frequency shift with short *T*_2_ *(*T* _2_ *_*short*_), and long *T*_2_ *(*T* _2_ *_*long*_), respectively. After the fitting, the voxel-wised ratio between *s*_0,*short*_ and the total proton density (*s*_0,*short*_ + *s*_0,*long*_) was reported as ultra-short T_2_ fraction map, and T_2_*_*short*_ and T_2_*_*long*_ were reported as ultra-short T_2_ value and long T_2_ value maps, correspondingly. All maps were registered to a standard brain atlas (MNI-152) for better visualization and then were averaged across subjects for statistical analysis. The ultra-short T_2_ fractions in total WM and total GM were reported individually and as mean across subjects. A two-sample t-test was used to test whether the ultra-short T_2_ fractions were statistically different between GM and WM. In addition, the ultra-short T_2_ fractions were quantified based on selected regions of interest (ROIs), including cingulum, corona radiata (CR), internal capsule (IC), corpus callosum (CC), external capsule, fornix, and sagittal stratum.

**Figure 2.**
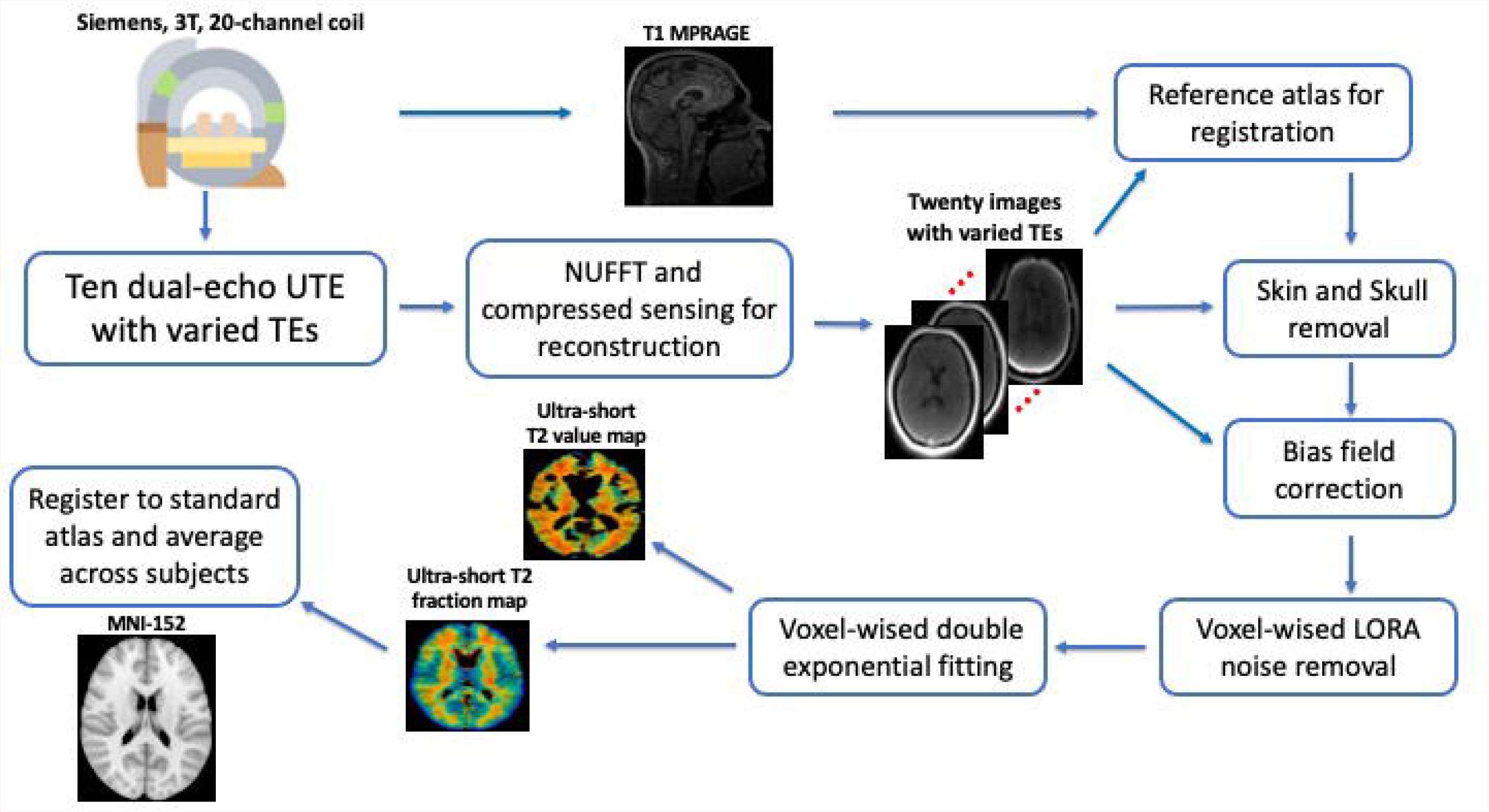
Workflow from data acquisition, image reconstruction to post-processing. NUFFT: nonuniform fast Fourier transform.

A phantom and one volunteer were scanned using the dual-echo 3D rosette UTE sequence and a 3D radial UTE sequence with golden angle acquisition (39) to compare the image quality after reconstruction. For this comparison, most of the imaging parameters were identical or closest possible values between the two sequences, which were: number of spokes (petals)=36100, TE_rosette_=20 *μs*, TE_radial_=30 *μs*, TR=7 ms, flip angle=7-degree, FOV=240×240×240 mm^3^, matrix size=128×128×128, readout dwell time=10 *μs*, and RF pulse duration=10 *μs*. For the dual-echo 3D rosette UTE sequence,105 samples were acquired per petal per TE, while 128 samples were acquired per spoke for 3D radial UTE. Two reconstruction methods were performed: compressed sensing (as described above) and regular regridding applying a density compensated adjoint NUFFT. The reconstruction processing steps were kept identical for the data acquired by both the dual-echo 3D rosette and radial UTE sequences.

## Results

Figure 3 (phantom) and Figure 4 (*in vivo*) compare the dual-echo 3D rosette UTE sequence and the 3D radial UTE sequence. All the reconstructed images were normalized to have signal intensity ranging from 0 to 1. The standard regridding reconstruction method resulted in ‘ringing’ artifacts for both acquisition methods (as shown in Figure 3B, 3D, 3F and Figure 4B, 4D, 4F), which had been minimized by the compressed sensing reconstruction method (as shown in Figure 3A, 3C, 3E and Figure 4A, 4C, 4E, correspondingly). The reconstructed image quality by the dual-echo 3D rosette UTE sequence was overall better than the 3D radial UTE, in terms of SNR (as shown in Figure 3A vs. 3E, and Figure 4A vs. 4E), and the ability to capture additional structural information (white dots as shown in Figure 3A, but not shown in Figure 3E). In addition, the dual-echo 3D rosette sequence detected the rapid decaying signals originating from the foam pad used for positioning (Supporting Information Figure S1). However, the foam pad was not identified by the 3D radial sequence.

**Figure 3.**
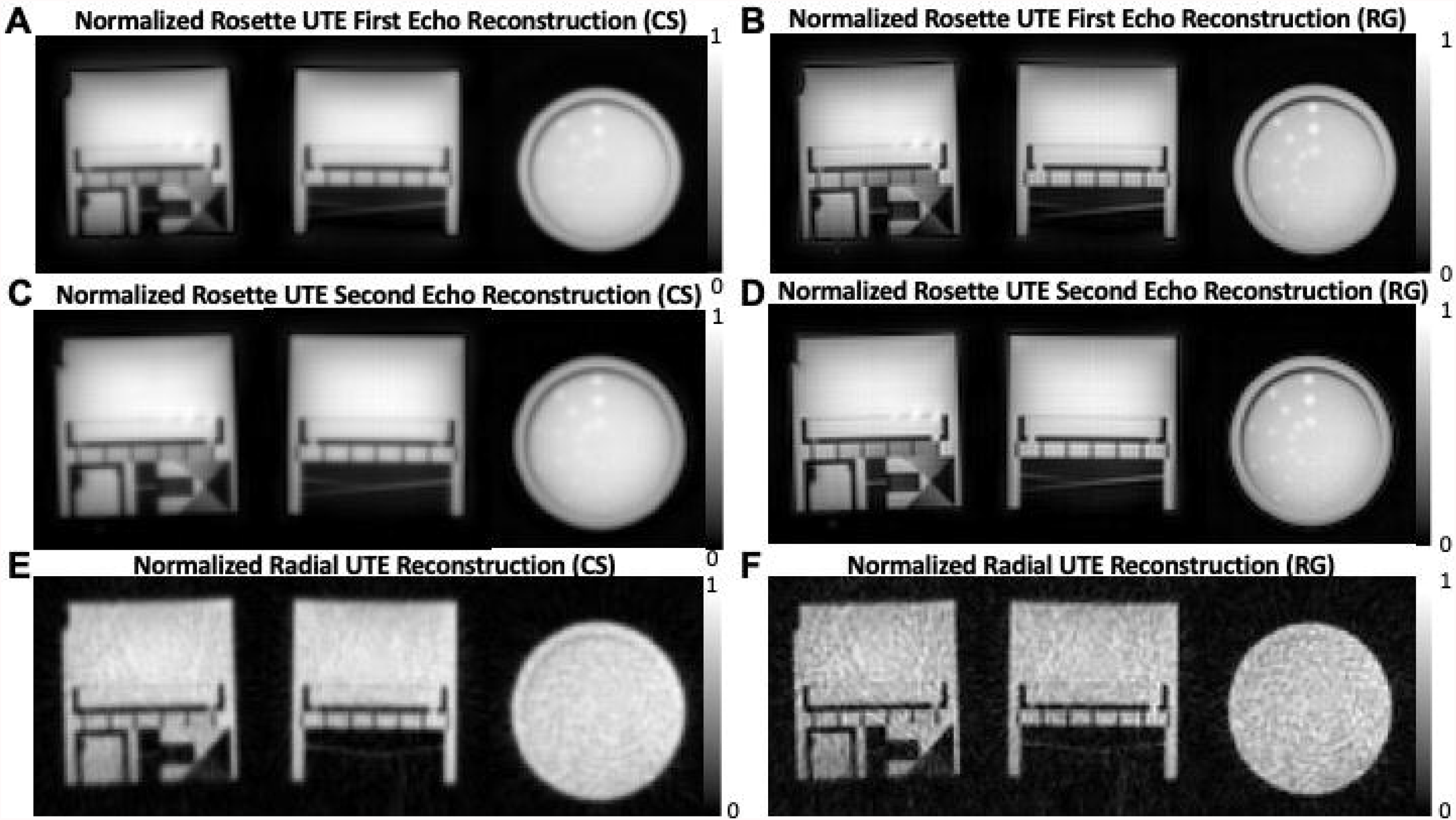
Phantom performance comparison between UTE sequences with 3D dual-echo rosette trajectory and 3D radial golden angle trajectory. All the images were normalized to have signal intensity in the range of 0 to 1. A, C, E) Reconstructed images based on compressed sensing (CS) technique from data sampled by A) the first echo of rosette trajectory; C) the second echo of rosette trajectory; E) radial trajectory. B, D, F) Reconstructed images based on regridding (RG) technique from data sampled by B) the first echo of rosette trajectory; D) the second echo of rosette trajectory; F) radial trajectory.

**Figure 4.**
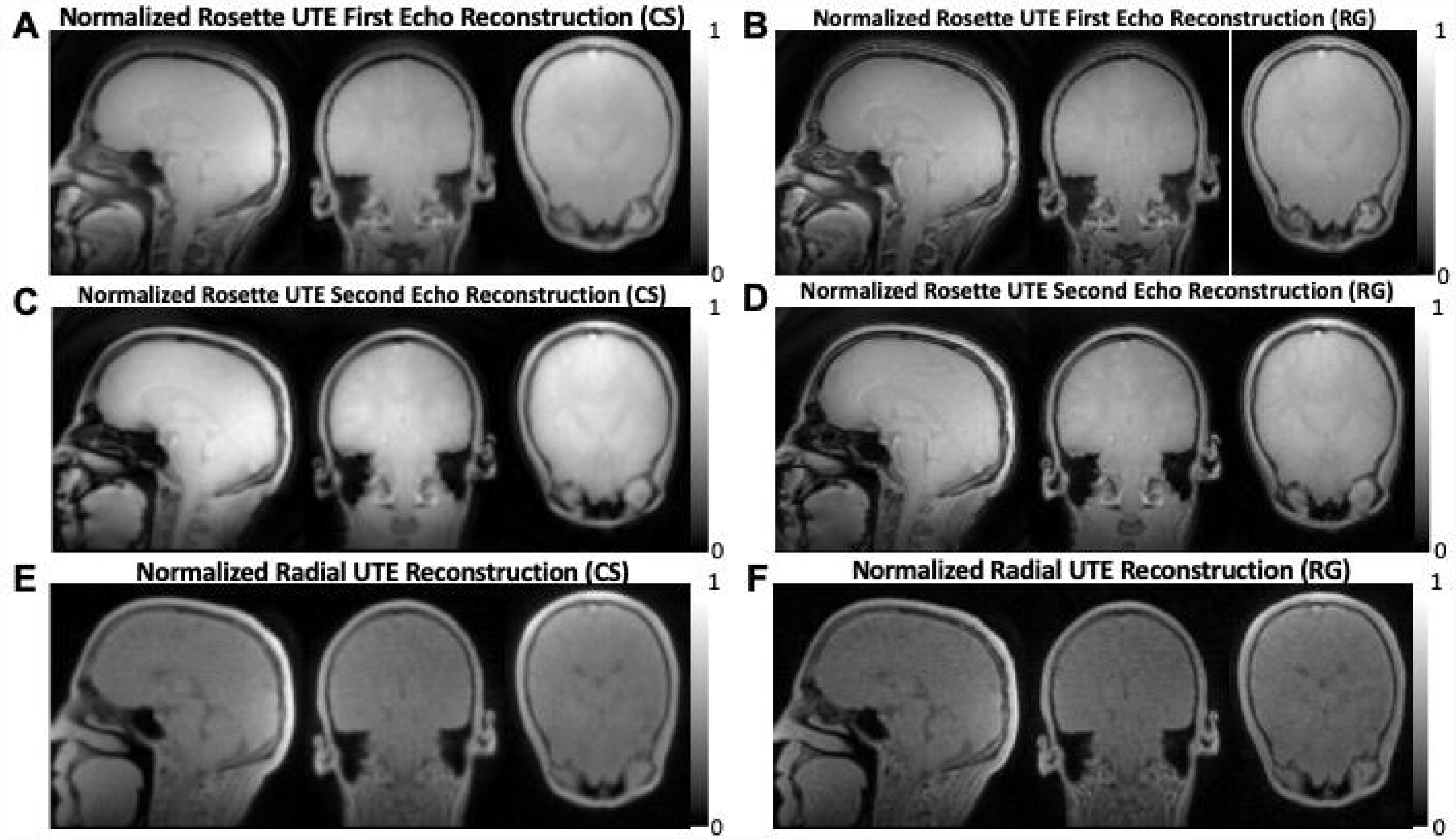
*In vivo* performance comparison between UTE sequences with 3D dual-echo rosette trajectory and 3D radial golden angle. All the images were normalized to have signal intensity in the range of 0 to 1. A, C, E) Reconstructed images based on compressed sensing (CS) technique from data sampled by A) the first echo of rosette trajectory; C) the second echo of rosette trajectory; E) radial trajectory. B, D, F) Reconstructed images based on regridding (RG) technique from data sampled by B) the first echo of rosette trajectory; D) the second echo of rosette trajectory; F) radial trajectory.

In Figure 5A, brain image slices from a volunteer are shown for five representative TEs, 20 μs, 100 μs, 2.12 ms, 2.4 ms, and 3.6 ms. There was little or no contrast between WM and GM at the minimum ultra-short TE (TE=20 μs), whereas the contrast between WM and GM increased at longer TE.

**Figure 5.**
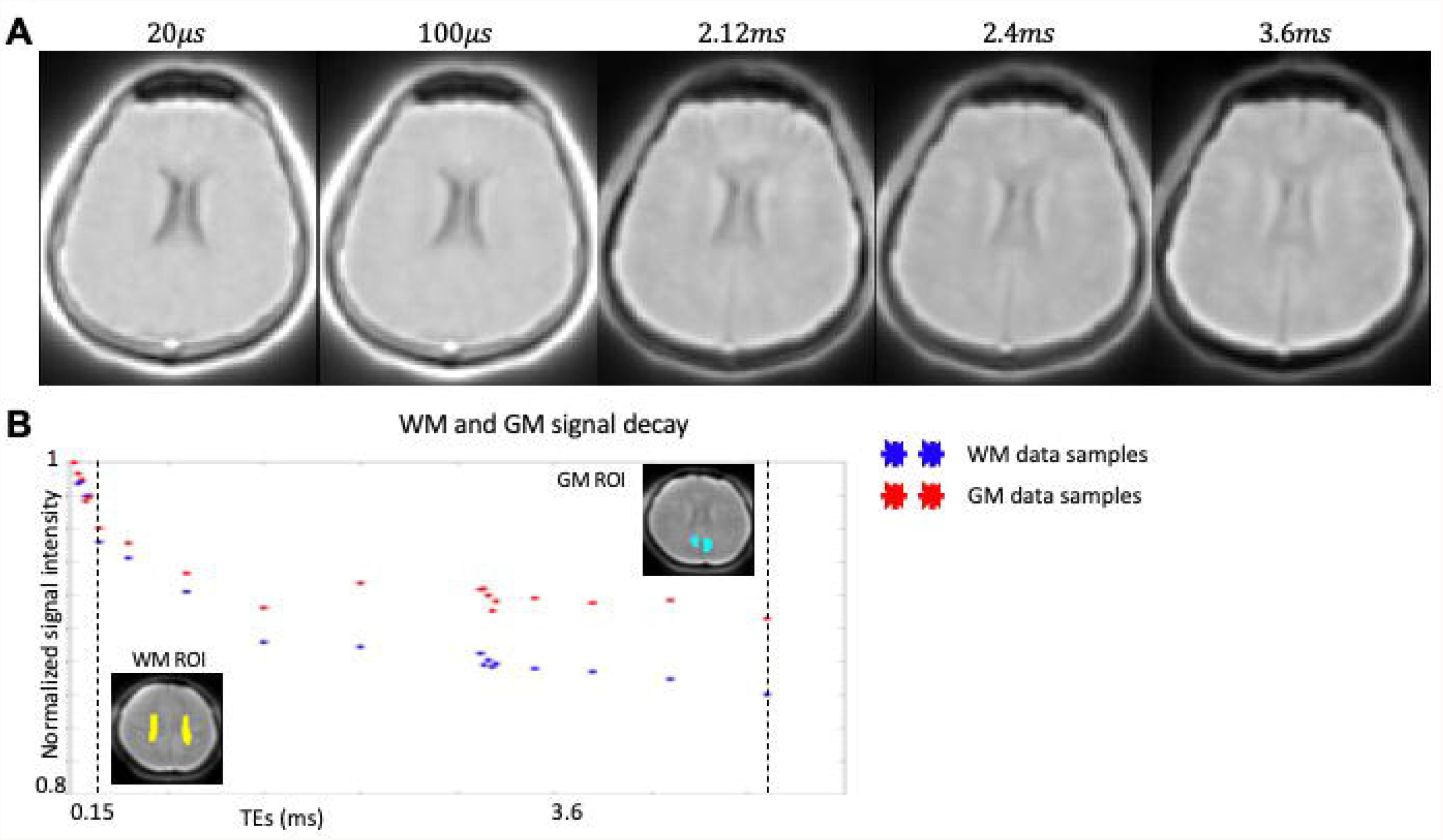
A) Five brain image slices with 20 μs, 100 μs, 2.1 ms, 2.4 ms, and 3.6 ms TEs from a volunteer are shown. B) Signal decay curves from selected regions of interest (ROIs), which are identified as white matter (WM) ROI (left bottom corner) and gray matter (GM) ROI (right upper corner). All signals were normalized based on the shortest TE (∼20 μs). The stars showed the distribution of signals with different TEs from WM ROI (blue) and GM ROI (red). Black dash lines were used to identify the ultra-short TEs region (below 0.15 ms).

Figures 5B shows the differences in signal decay between WM ROI (blue) and GM ROI (red). The signal intensities with different TEs were normalized based on the detected signal at the shortest TE of 20 μs. WM and GM signal decay curves showed fast signal drop at ultra-short TE period (below 0.15 ms), which was about 5% in selected WM ROI and about 4% in GM ROI. The signal curve of WM ROI continued to drop (about 15% at TE=3.6ms), while the signal curve of GM ROI had a slower decay at longer TEs (about 10% at TE=3.6ms).

Table 1 summarizes the ultra-short T_2_ components (uT_2_) fraction in total GM and total WM individually and as mean across subjects. All subjects and the average indicated a significantly higher uT_2_ fraction in WM than GM (*P*<0.0001 for all subjects and the mean). The average WM uT_2_ fraction across subjects was 10.9%±1.9%, and the average GM uT_2_ fraction across subjects was 5.7%±2.4%.

**Table 1.** Mean and standard deviation of the ultra-short T_2_ components (myelin) fraction in the grey matter (GM) and white matter (WM) Individual and mean across subjects ultra-short T_2_ fractions in total white matter (WM) and total grey matter (GM). All values of ultra-short T_2_ fractions are shown as mean±standard deviation. A two-sample t-test was used to test whether the ultra-short T_2_ fractions were statistically different between GM and WM, which *P*-values were calculated.

| Subject number | Total GM | Total WM | *P*-values |
| --- | --- | --- | --- |
| 1 | 7.5%±1.1% | 11.6%±0.9% | *P*<0.0001 |
| 2 | 3.7%±1.4% | 9.9%±1.3% | *P*<0.0001 |
| 3 | 3.9%±2.1% | 9.1%±1.0% | *P*<0.0001 |
| 4 | 8.7%±1.4% | 13.8%±1.2% | *P*<0.0001 |
| 5 | 5.4%±1.6% | 10.5%±1.1% | *P*<0.0001 |
| Total | 5.7%±2.4% | 10.9%±1.9% | *P*<0.0001 |

Figure 6 shows the mean uT_2_ fraction map (Figure 6A), the mean ultra-short T_2_ value map (Figure 6B), and the mean long T_2_ value map (Figure 6C) in MNI-152 space. The uT_2_ fraction map (Figure 6A) indicated a generally homogeneous uT_2_ fraction among voxels in WM, which was higher than the uT_2_ fraction among voxels in GM. The ultra-short T_2_ value in WM was around 0.15 ms, and the ultra-short T_2_ value in GM was slightly faster than WM (0.1 ms). The long T_2_ components T_2_ value map indicated an increased T_2_ value across GM and the CSF compared to the WM, which is in line with previous reports (37,40). (The uT_2_ fraction maps and ultra-short T_2_ value maps from two subjects are shown in Supporting Information Figure S2).

**Figure 6.**
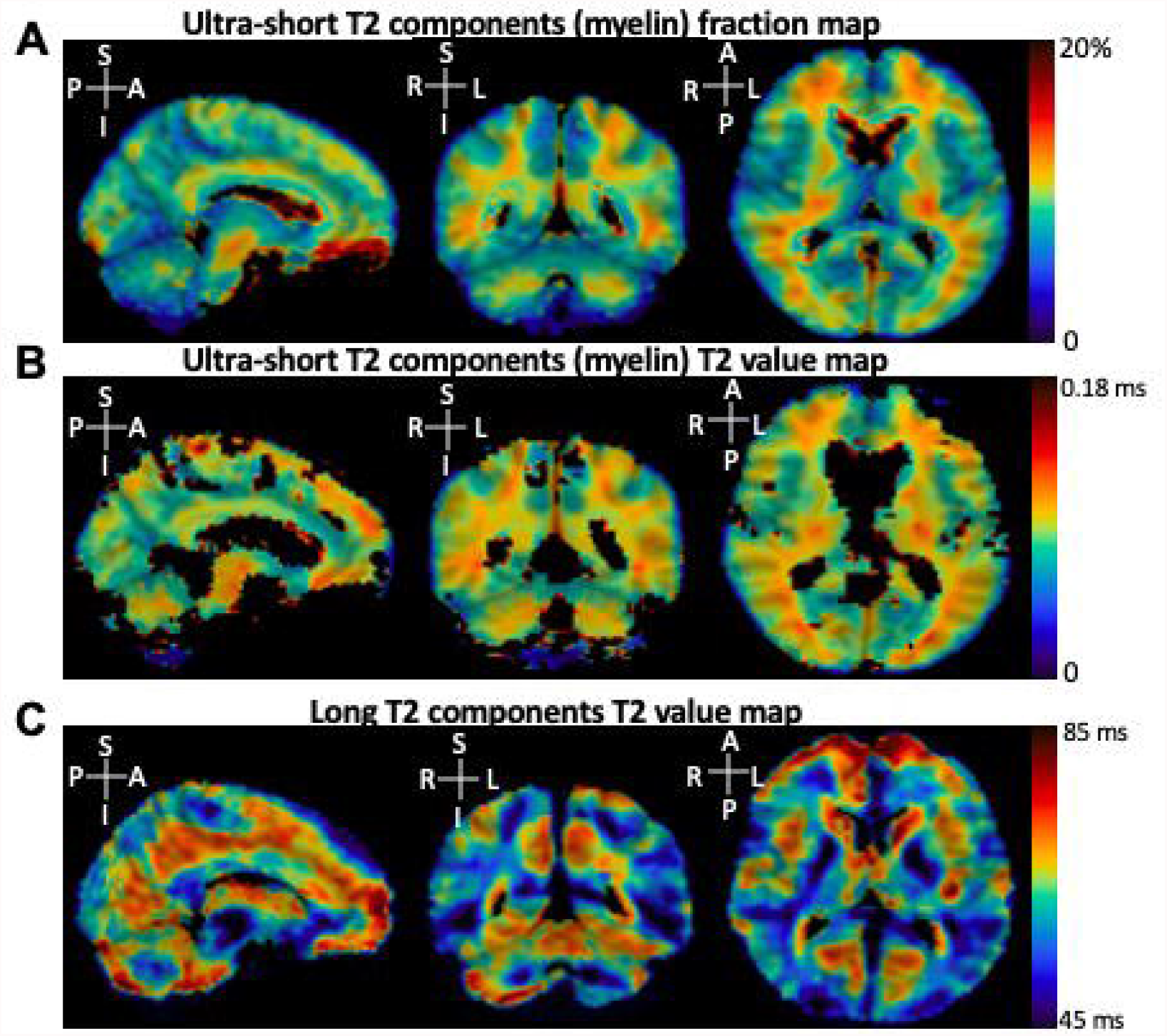
The results of dual-exponential fitting: A) The mean ultra-short T_2_ components (uT_2_) fraction map; B) The mean uT_2_ value map; C) The long T_2_ value map. A: anterior, P: posterior, L: left, R: right, S: superior, I: inferior. Please note that some voxels were black in Figure 4B (primarily located in cerebrospinal fluid and grey matter), caused by high uT2 values reaching the top threshold of the color bar.

Figure 7 summarizes the mean and standard deviation (SD) of the uT_2_ fraction among different ROIs across five subjects (cingulum, corona radiata, internal capsule, corpus callosum, external capsule, fornix, and sagittal stratum). All the ROIs were identified as WM-rich regions and reported about 10% as the uT_2_ fraction. The lowest reported ROI-based uT_2_ fraction value, 9.4%±1.0%, was in the corpus callosum. The highest reported ROI-based uT_2_ fraction value, 12.2%±1.6%, was in the fornix.

**Figure 7.**
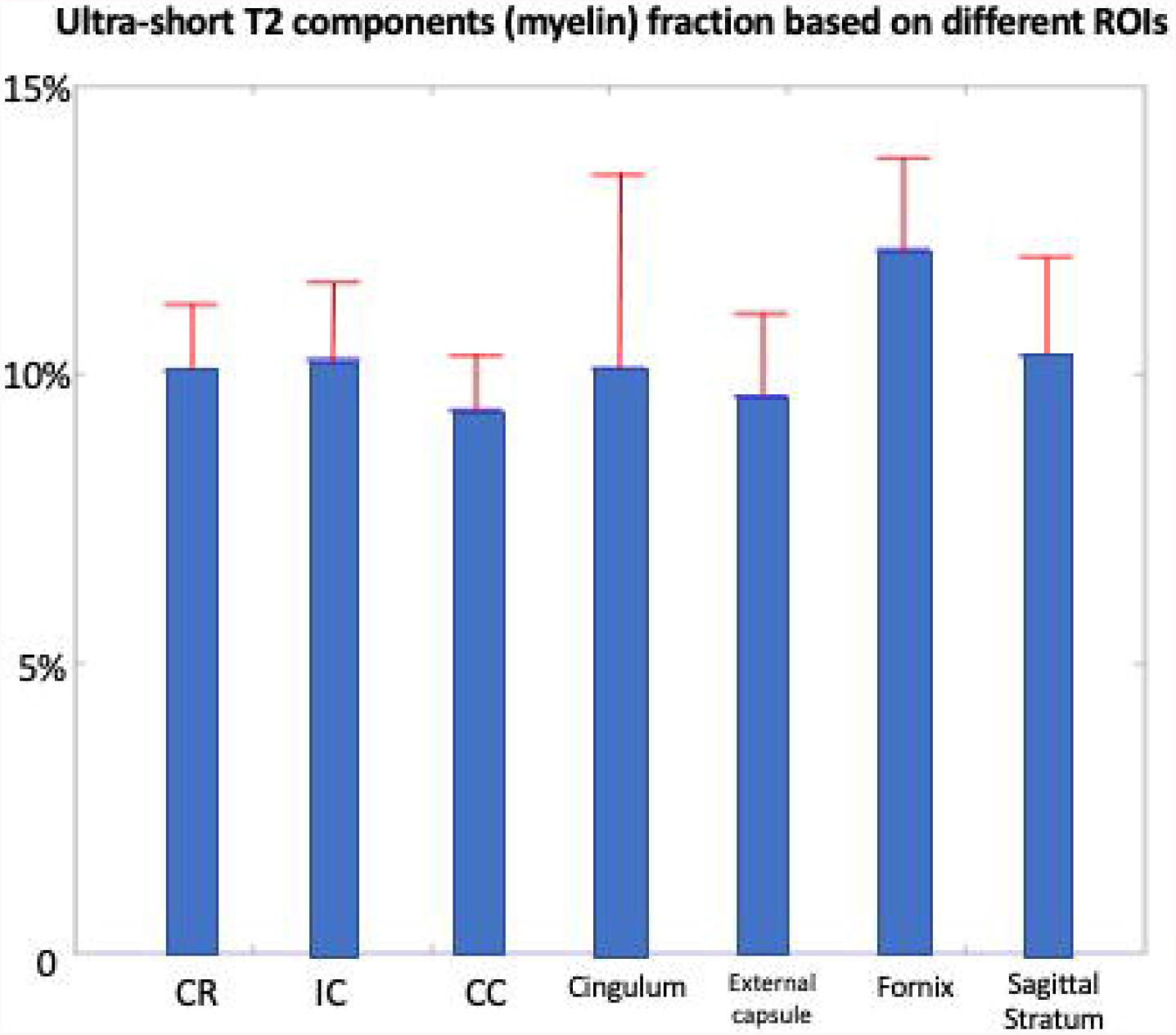
The mean (blue columns) and standard deviation (SD) (red) of ultra-short T_2_ components (uT_2_) fraction among different regions of interest (ROIs) across subjects, quantified based on the five individual uT_2_ fraction maps. CR: corona radiata, IC: internal capsule, CC: corpus callosum.

## Discussion

In this study, a novel 3D dual-echo UTE sequence based on rosette k-space trajectory was developed. The advantages of this novel design included: 1) increased sampling density in the outer k-space; 2) shorter TE due to the gradient design; 3) a smooth transition (zero-delay) between the two echoes of dual-echo acquisition. As a result, a statistically significantly higher uT_2_ fraction value was found in WM compared to GM. In addition, the uT_2_ fraction value was homogeneously distributed among WM voxels.

The achievement of the shortest TE in UTE sequences is limited by the system hardware, not only by the dead time (13) but also by the slew rate limitation if the initial gradient amplitude is high in any direction. In this specific 3D rosette k-space design, the initial gradient amplitude was zero and gradually increased to overcome the slew rate limitation in all acquisitions. The strategy was to gradually increase the *ω*_l_ and *ω*_2_ to the designed value of 1.611 kHz at the beginning of each TR. With this strategy, the maximum slew rate was about 140 mT/m/ms for all three directions, separately. In addition, the initial sampled data still followed the originally designed rosette trajectory, which was used for image reconstruction. This resulted in an over-sampling at the center k-space (about 20% samples located in radii of k-space smaller than 0.25*Kmax*), which could provide an additional signal-to-noise ratio (SNR). Similar to ZTE applications, this study used a small flip angle (7 degrees). Although large flip angles are allowed in UTE acquisition, a small flip angle could minimize the T_1_ influence on the detected signals (41).

The volume of 3D k-space is in proportion to the cube of the radius of the acquired k-space. The volume of radii larger than 0.75 *Kmax* (considered as peripheral k-space) occupied about 58% (= (1^3^-0.75^3^)/1^3^) of the entire volume of 3D k-space with the radius of *Kmax*. However, the volume of radii smaller than 0.25 *Kmax* (considered as center k-space) only occupied about 2% (= 0.25^3^/1^3^) of the entire volume of k-space. This resulted in a much larger peripheral k-space than the center k-space volume in 3D cases. The conventional radial acquisition, which samples uniformly along with the 1D radius of k-space (25% of samples at center k-space, and 25% of samples at peripheral k-space), may not be efficient at the large radii of 3D k-space. Under specific circumstance, e.g., 80% undersampling number of spokes, the reconstructed images acquired by the 3D radial pattern resulted in noise amplification and the loss of structural details. In addition to the insufficient samples at the large radii of 3D k-space, the nonoptimal imaging parameters (e.g., insufficient number of spokes, long TR), which were set to match 3D dual-echo rosette UTE, also contributed to the degraded performance of 3D radial acquisition.

The newly proposed 3D rosette trajectory has increased samples at larger radii of k-space, which provides greater sampling density in peripheral k-space. The advantages of the newly proposed 3D rosette trajectory (i.e., trajectory curvature and k-space coverage) may have the potential for further acceleration by acquiring fewer petals without losing image quality. As stated in the Theory section, the number of petals (spokes) used to compare the dual-echo 3D rosette and the 3D radial trajectories is an 80% undersampling pattern. However, the reconstructed images using CS technique acquired by the dual-echo 3D rosette trajectory did not suffer from significant artifacts and did not lose structural information compared to the images acquired by the 3D radial scan. In addition, although the regridding method resulted in sharper images, it suffers ringing artifact due the undsampling. In this study, in order to minimize the ringining artifact, CS method was used with a cost of losing fine details. Adapting this dual-echo 3D rosette trajectory to keep the image quality with an undersampling strategy may lead to future directions. For example, with additional undersampling factor=2 (40%) or 4 (20%), the total scan time of the ten dual-echo UTE sequences can be shortened to between 21 and 10 minutes, which could be acceptable in clinical settings (reconstructed images with undersampling factor=2, which means number of petals=18050 are shown in Supporting Information Figure S3).

Instead of combining IR suppression and dual-echo subtraction, which provides myelin maps with arbitrary signal intensities (7), dual-exponential complex fitting was performed in this study, offering uT_2_ fraction and uT_2_ value maps. Although the quantitative analysis of myelin by the uT_2_ fraction and value maps still needs validation (7), this method showed a potential way to compare across subjects. The setback of this method was the need for a multiple TEs acquisition, which increased total scan time. However, with the dual-echo rosette sequence proposed in this study, twenty images with different TEs could be acquired in a reasonable total scan time. A clinically acceptable scan time can be achieved with a further reduction of acquisition duration with an undersampling strategy. On the other hand, when the dual-echo subtraction method has been applied to the first and second TEs of the dual-echo 3D rosette to remove the long T_2_ components, this will create mono-exponential signal decay predominantly originating from the short-T_2_ components (Supporting Information Figure S4).

Overall, the results of uT_2_ components quantification, acquired by the dual-echo 3D rosette sequence, agree with previous publications with 3D radial or 3D cones trajectories (7,42). Firstly, little or no contrast at short TEs (i.e., below 100 *μs*) was achieved without IR suppression. Secondly, the contrast between WM and GM became higher at longer TEs, as the uT_2_ components in WM started to decay. Thirdly, the uT_2_ components were mainly distributed in WM voxels, resulting in a generally higher uT_2_ fraction value in WM compared to GM. Additionally, the uT_2_ fraction values based on ROIs aligned with previous publications (43-45), around 10% to 12% in most WM-rich brain regions.

One disadvantage of the 3D rosette k-space pattern is that more samples per acquisition were needed (1.5 times samples compared to radial). In addition, the long readout duration of this rosette trajectory (2.1 ms for dual-echo acquisition, which means 1.05 ms for the first echo acquisition) may suffer image blurring caused by the fast signal decay (6). A previous study has shown that the ideal sampling duration is 0.81*T_2_ of the imaging tissue for radial UTE (46). However, such a short sampling duration would result in spatial resolution loss (6). Therefore, typically, the sampling duration of two to four times T_2_ of the imaging tissue was used (6). For the myelin imaging applications, the sampling duration was usually set to 0.3 to 1 ms (1,23,47). One of the significant issues of model-based processing is to fit a large number of parameters, long and ultra-short signal decay rates, and respective frequency shifts. This results in complications with the degradation of the accuracy and precision of the parameter estimations. This study used magnitude data for the complex model. Although this approach resulted in stable frequency differences between WM and GM, absolute frequency offsets could not be estimated. On the other hand, we attempted to use the complex data for the complex model (37). However, the estimations hit the boundary conditions and resulted in unstable and hence inhomogeneous estimates of parameters in 2 out of 5 subjects (data was not shown). This might be due to the relatively long acquisition duration for both echoes (2.1 ms), propagation of phase errors due to eddy currents, gradient delays, and breathing. One can overcome these limitations using field cameras (48). Alternatively, the gradient impulse response function can be calculated to mimic the behavior of the gradient chain and then use it for the image reconstruction retrospectively (49). In particular, the potential use of these methods may overcome the limitations of the usefulness of the second echoes of the dual-echo 3D rosette technique.

Recent publications showed that the brain iron content led to inaccurate results of the myelin water fraction (MWF) (about 26%-28% decrease after iron extraction by reductive dissolution of brain slice samples) by myelin water imaging technique (MWI) (50). Although 3D UTE rather than MWI was used in this study, the influence of iron content on the results was not ruled out. Myelin concentrations could be higher than expected since the iron content decreased the MR signal, especially within substantia nigra (51), putamen (52), and globus pallidus (53) regions, which have potential age-related iron accumulation. Those regions had high signal intensities in our reconstructed images of the novel dual-echo 3D rosette acquisition with the shortest TE (∼ 20 *μs*). A new model for fitting could regress the iron effect out (54). In addition, quantitative susceptibility mapping (55) may be applied for iron measurement since the proposed sequence allows a dual-echo acquisition.

There are some other limitations of this study. Firstly, the sample pool of five healthy volunteers was limited. Due to its nature as proof of concept, the sensitivity of the novel 3D rosette trajectory UTE sequence, and/or the dual-exponential model fitting method to detect diseased states (e.g., demyelination) is undetermined. A prospective study is required to address this limitation. Secondly, other macromolecules with similar uT_2_ values may contribute to the detected signals (56). For future studies, patients (and/or animals) with different stages of myelination diseases will be recruited to test the performance of this novel rosette k-space trajectory.

In conclusion, this study proposed a novel dual-echo 3D rosette k-space trajectory, specifically for UTE applications. The higher uT_2_ fraction value in WM compared to GM demonstrated the ability of this sequence to capture rapidly decayed signals. In addition, the fitting based on the dual-exponential model provided quantitative results of the uT_2_ fraction, which could be used for myelination assessment in the future.

## Acknowledgement

Data acquisition was supported in part by NIH grant S10 OD012336. In addition, this project was supported by an award from the Ralph W. and Grace M. Showalter Research Trust.

The authors are grateful to Prof. Mehmet Ali Deveci from Koc Univesity, Istanbul Turkey for the clinical discussion.

## Notes

### Competing Interest Statement

The authors have declared no competing interest.

### Summary of Updates

The content of the manuscript has been modified to address these, including 1) a comparison between the dual-echo 3D rosette and the conventional 3D golden angle radial UTE; 2)reconstruction results from regular regridding; 3) reconstructed images sampled by undersampling dual-echo 3D rosette patterns to demonstrate the prospective acceleration; 4) ultra-short T2 fraction and value maps from two representative individuals and 5) potential use of the second echo.

