## Supplemental Figures for "Myelin Imaging Using Dual-echo 3D Ultra-short Echo Time MRI with Rosette k-Space Pattern"

Supporting Information Figure S1

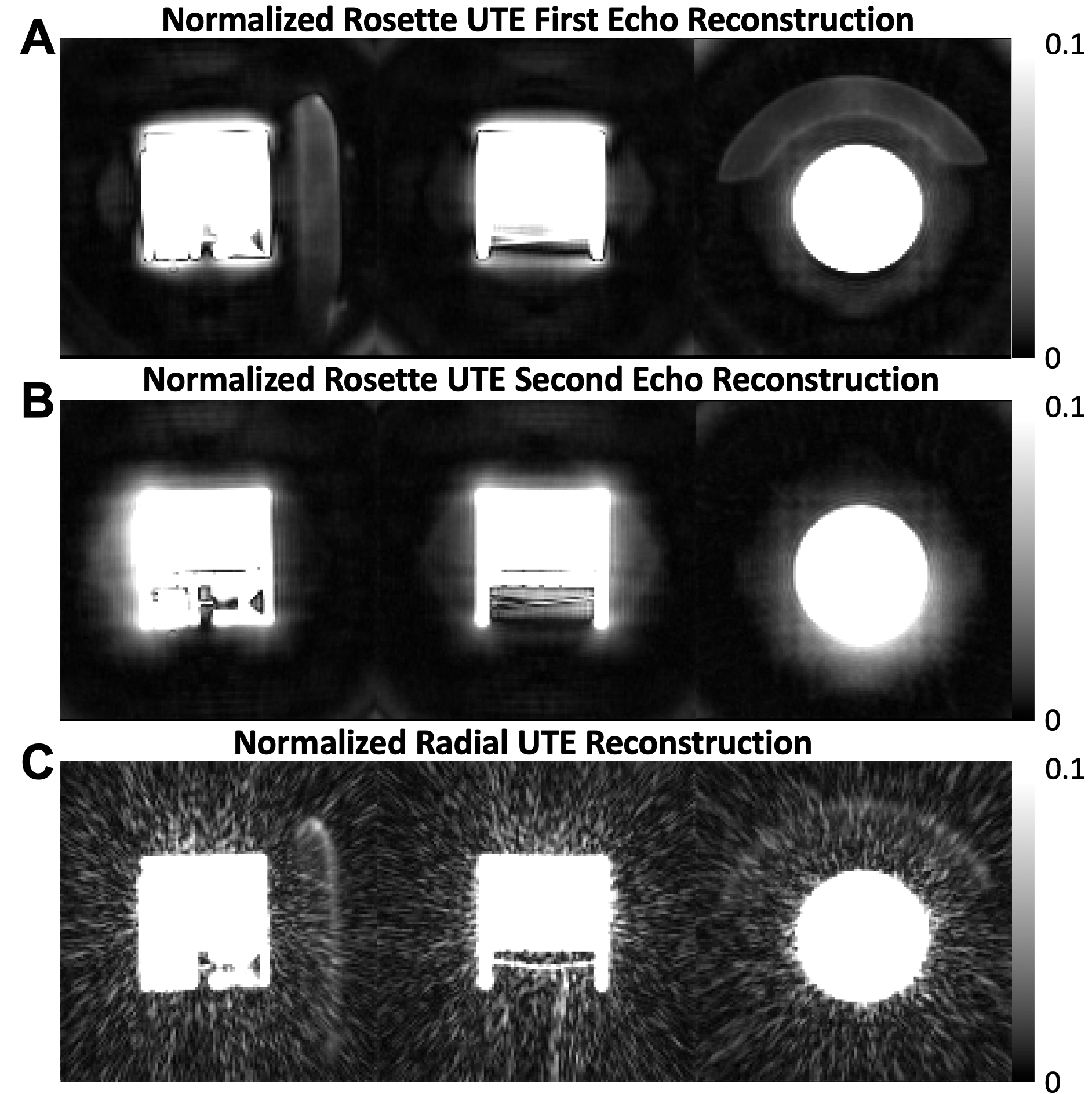

Supporting Information Figure S1 Caption

Performance comparison between UTE sequences with dual-echo 3D rosette trajectory and 3D radial golden angle trajectory based on the phantom scan. All the images were reconstructed based on the compressed sensing technique described in the Methods section and were normalized to have signal intensity in the range of 0 to 1. A colorbar with a reduced threshold (0 to 0.1) was used to highlight the foam pad used for positioning. The dual-echo 3D rosette sequence with ultra-short TE (20 μs) (A) detected the rapid decaying signals originating from the foam pad. However, the foam pad was not identified by the dial-echo 3D rosette sequence with longer TE (2.12 ms) (B) or by the 3D radial sequence with ultra-short TE (30 μs) (C).

Supporting Information Figure S2

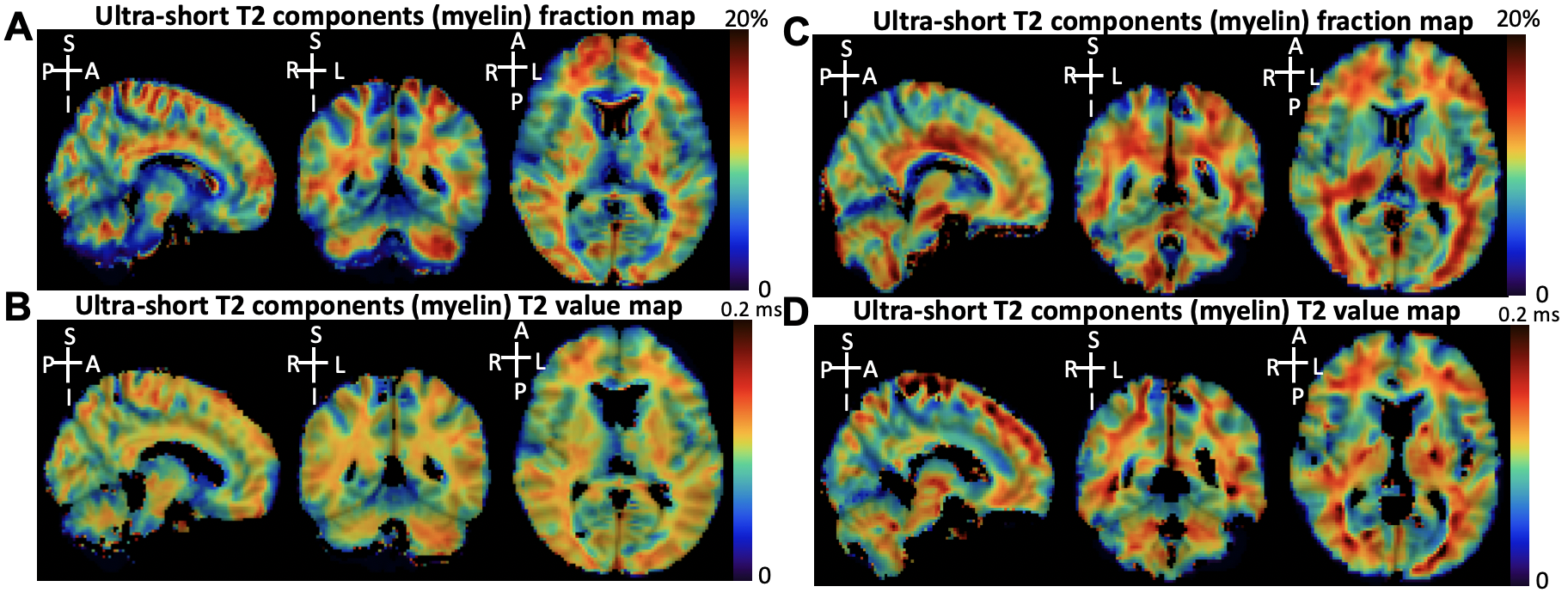

Supporting Information Figure S2 Caption

Two representative subjects’ results of ultra-short T_2_ components (uT_2_) fraction maps and ultra-short T_2_ value maps are illustrated. A, B) uT_2_ fraction maps (A) and ultra-short T_2_ value maps (B) from one volunteer. C, D) uT_2_ fraction maps (C) and ultra-short T_2_ value maps (D) from another volunteer (different volunteer). A: anterior, P: posterior, L: left, R: right, S: superior, I: inferior.

Supporting Information Figure S3

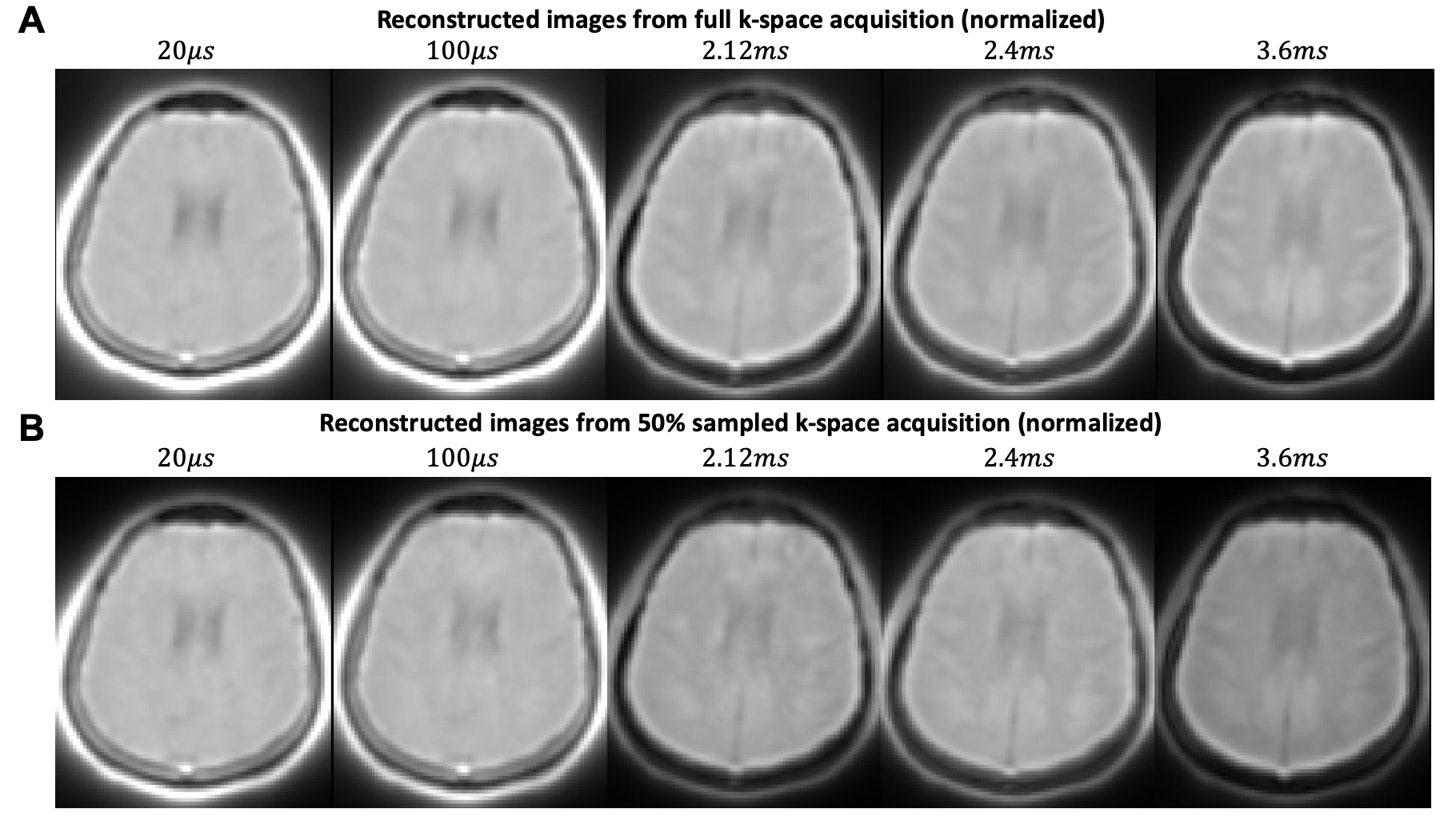

Supporting Information Figure S3 Caption

Reconstructed brain image slices based on compressed sensing technique for the dual-echo 3D rosette with TEs of 20 μs, 100 μs, 2.12 ms, 2.4 ms, and 3.6 ms from a volunteer are shown. All the image slices were normalized to have signal intensity in the range of 0 to 1. A) Reconstruction results from full k-space acquisition (number of petals=36100). B) Reconstruction results from 50% k-space acquisition (undersampling factor=2, number of petals=18050).

Supporting Information Figure S4

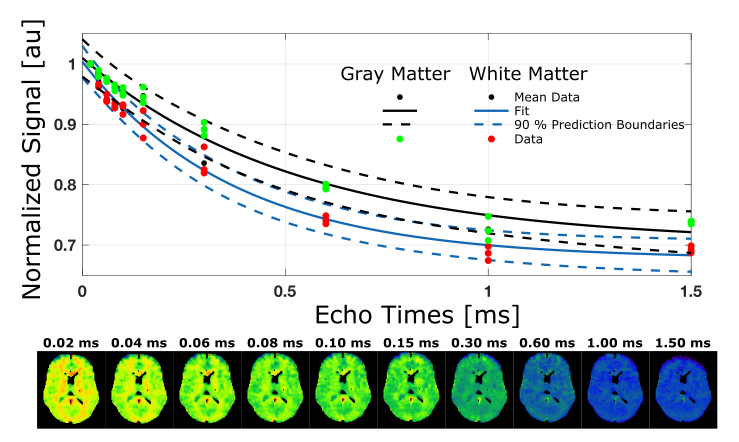

Supporting Information Figure S4 Caption

The potential use of the Second TE. Direct subtraction (first minus second TE image, i.e., TE_1_=20 ms minus TE_2_=2120 ms) contrasts white and grey matter. Increasing TE leads to a decrease in the distinction between white matter and gray matter. This demonstrates uT_2_ component predominantly originates from white matter. Estimated uT_2_ values of mono-exponential fittings are 526 μs for the gray matter and 375 μs for the white matter.
